# Inbreeding and gallbladder cancer risk: Homozygosity associations adjusted for indigenous American ancestry, BMI and genetic risk of gallstone disease

**DOI:** 10.1101/2024.04.22.590517

**Authors:** Francisco Ceballos, Felix Boekstegers, Dominique Scherer, Carol Barahona Ponce, Katherine Marcelain, Valentina Gárate-Calderón, Melanie Waldenberger, Erik Morales, Armando Rojas, César Munoz, Javier Retamales, Gonzalo de Toro, Allan Vera Kortmann, Olga Barajas, María Teresa Rivera, Analía Cortés, Denisse Loader, Javiera Saavedra, Lorena Gutiérrez, Alejandro Ortega, Maria Enriqueta Bertrán, Leonardo Bartolotti, Fernando Gabler, Mónica Campos, Juan Alvarado, Fabricio Moisán, Loreto Spencer, Bruno Nervi, Daniel Carvajal-Hausdorf, Héctor Losada, Mauricio Almau, Plinio Fernández, Jordi Olloquequi, Francisco Rothhammer, Justo Lorenzo Bermejo

## Abstract

Latin Americans have a rich genetic make-up that translates into heterogeneous fractions of the autosomal genome in runs of homozygosity (F_ROH_), and heterogeneous types and proportions of indigenous American ancestry. While autozygosity has been linked to several human diseases, very little is known about the relationship between inbreeding, genetic ancestry and cancer risk in Latin Americans.

Chile has one of the highest incidences of gallbladder cancer (GBC) in the world, and here we investigated the association between inbreeding, GBC, gallstone disease (GSD) and body mass index (BMI) in 4029 genetically admixed Chileans. We calculated individual F_ROH_ above 1.5 Mb and weighted polygenic risk scores for GSD, and applied multiple logistic regression to assess the association between homozygosity and GBC risk.

We found that homozygosity was due to a heterogeneous mixture of genetic drift and consanguinity in the study population. Although we found no association between homozygosity and overall GBC risk, we detected interactions between F_ROH_ and sex, age, and genetic risk of GSD on GBC risk. Specifically, the increase in GBC risk per 1% F_ROH_ was 19% in men (P-value = 0.002), 30% in those under 60 years of age (P-value = 0.001), and 12% in those with a genetic risk of GSD above the median (P-value = 0.01).

The present study highlights the complex interplay between inbreeding, genetic ancestry and genetic risk of GSD in the development of GBC. The applied methodology and our findings underscore the importance of considering the population-specific genetic architecture, along with sex- and age specific-effects, when investigating the genetic basis of complex traits in Latin Americans.

## Introduction

Gallbladder cancer (GBC) remains an aggressive disease with very limited treatment options and a lack of reliable markers for early detection (1, 2). As of 2020, the incidence of GBC is projected to increase by 75% by 2040, underscoring the urgency to characterize the factors that contribute to GBC development (3). Currently, the best predictors of GBC risk include the presence of gallstones, as well as age and sex, with women being more susceptible to the disease.

Large differences in the incidence and mortality of GBC are observed in different populations and geographic regions, challenging our understanding of GBC etiology (4, 5). The highest incidences have been reported in Bolivia (especially around Lake Titicaca), Chile (especially in the southern regions), Peru (especially in the city of Trujillo), Japan, northern India, and New Mexico (1, 4). This geographical clustering suggests a possible link between GBC development and ancestry, particularly in individuals with indigenous Asian and American roots, which may have a genetic, cultural or mixed origin.

Among these clusters, Chile stands out as the country with the highest GBC incidence, with approximately 27.3 cases per 100,000 individuals (1). Even within Chile, GBC incidence shows considerable heterogeneity, further highlighting the potential role of ancestry in disease susceptibility (6-8). The relatively simple distribution of ancestry components in Chile facilitates the study of the genetic basis of GBC. The African contribution to the Chilean genome is limited (<3% on average), and the proportion of European ancestry is particularly high in the central metropolitan region (9-11). The indigenous American ancestry can be broadly divided into two main components: Aymara-Quechua ancestry in northern Chile, and Mapuche-Huilliche ancestry in the south. Notably, in contrast to Aymara-Quechua ancestry, each 1% increase in the individual proportion of Mapuche-Huilliche ancestry was associated with a 2% increased risk of developing GBC and a 3.7% higher GBC mortality (5). Consistent with this association, the prevalence of GBC is about 20 times higher in Argentina’s Andean region than in the rest of the country, indicating a possible contribution of indigenous American ancestry to GBC susceptibility in this region as well (3). Other GBC risk factors such as gallstone disease (GSD), elevated body mass index (BMI), low socioeconomic status, and lifestyle in general could confound the association between indigenous American ancestry and GBC risk, but the results of a recent study suggest a putatively causal effect of Mapuche-Huilliche ancestry on GBC development (12).

Genomic homozygosity, quantified by Runs of Homozygosity (ROH), i.e. contiguous stretches of homozygous alleles in identical-by-descent status, reflects the demographic history of both individuals and populations, and has been shown to influence several complex traits (13). Large studies have found associations between the fraction of the genome in ROH (F_ROH_) and a wide range of phenotypes, including height, BMI, diabetes, heart disease, and subcutaneous adipose tissue (14, 15). However, most published studies on the effects of inbreeding on human diseases, particularly cancer, have shown inconsistent results (13). Some of the reasons for this inconsistency are small sample sizes, limited F_ROH_ variability in the European outbred populations in which most of these studies have been conducted, and the lack of a standardized procedure for ROH analysis. Indigenous American genomes exhibit long stretches of homozygosity, Latin Americans are highly heterogeneous in terms of individual burden of homozygosity, and Chileans have been found to have both high ROH burden and high F_ROH_ variability (13, 16); (17).

In this context, the study of populations with a recent history of genetic admixture, and a high and variable degree of inbreeding, provides a unique opportunity to explore the relationship between genetic factors and the occurrence of GBC. In this study, we investigate the impact of homozygosity, quantified by individual F_ROH_ above 1.5 Mb, on GBC risk in Chileans. By simultaneously considering individual type and proportion of indigenous American ancestry, BMI and genetic risk of GSD, we aim to elucidate the mechanisms underlying geographical clustering of GBC, and potentially uncover novel genetic markers for predicting individual GBC risk.

## Results

**Table 1** shows the main characteristics of the study participants, both overall and stratified by specific subgroups, including GBC patients, who made up 15.3% of the study population, GSD patients (23.3%), and individuals classified as overweight (BMI > 25 kg/m^2^), who made up 61.5% of the study participants. On average, GBC patients were more often female, older, less educated and had a higher proportion of indigenous Mapuche-Huilliche ancestry than the total study population, while differences in genetic risk of GSD (quantified by weighted polygenic risk scores) and FROH were rather small (overlapping interquartile ranges [IQR]).

**Table 1.** Main characteristics of the study participants summarized by absolute and relative frequencies for categorical variables, and by medians and interquartile ranges for continuous variables.

| Variable | Level | All participants<br>n=4029 (100%) |  | Gallbladder cancer patients<br>n=616 (15.3%) |  | Gallstone disease patients<br>n=933 (23.2%) |  | Overweight participants<br>N=2254 (61.5%) |  |
| --- | --- | --- | --- | --- | --- | --- | --- | --- | --- |
| Sex | Male | 1744 | 43.3% | 147 | 23.9% | 189 | 20.3% | 1045 | 46.4% |
|  | Female | 2284 | 56.7% | 469 | 76.1% | 744 | 79.7% | 1209 | 53.6% |
| Age | Continuous | 37 | 26-58 | 60 | 49-67 | 56 | 41-66 | 40 | 28-59 |
| Education | Primary & informal schooling | 515 | 12.7% | 276 | 45.3% | 302 | 32.4% | 285 | 12.6% |
|  | Secondary | 1749 | 43.4% | 207 | 33.6% | 284 | 30.4% | 968 | 42.9% |
|  | Technical | 156 | 3.9% | 27 | 4.4% | 50 | 5.3% | 75 | 3.4% |
|  | Postgraduate | 69 | 1.7% | 3 | 0.5% | 7 | 0.8% | 42 | 1.9% |
|  | University | 532 | 13.2% | 39 | 6.3% | 77 | 8.3% | 280 | 12.4% |
|  | Missing | 1008 | 25.0% | 64 | 10.4% | 213 | 22.8% | 604 | 26.8% |
| Ancestry group | Aymara-Quechua | 111 | 2.7% | 7 | 1.1% | 12 | 1.3% | 57 | 2.5% |
|  | Aymara-Quechua-European | 197 | 4.9% | 3 | 0.5% | 8 | 0.9% | 114 | 5.1% |
|  | European | 113 | 2.8% | 15 | 2.5% | 23 | 2.5% | 51 | 2.3% |
|  | Mapuche-Huilliche | 82 | 2.1% | 39 | 6.3% | 39 | 4.2% | 48 | 2.1% |
|  | Mapuche-Huilliche-European | 1885 | 46.8% | 341 | 55.4% | 516 | 55.3% | 1069 | 47.4% |
|  | Other admixture | 1641 | 40.7% | 211 | 34.3% | 335 | 35.9% | 915 | 40.6% |
| Genetic risk of gallstone disease | Continuous | 0.45 | 0.38-0.53 | 0.45 | 0.39-0.54 | 0.47 | 0.39-0.54 | 0.45 | 0.38-0.54 |
| FROH | Continuous | 0.009 | 0.006-0.013 | 0.011 | 0.008-0.016 | 0.011 | 0.008-0.014 | 0.012 | 0.009-0.017 |

**Overweight participants:** Body mass index > 25 kg/m<sup>2</sup>, **Aymara-Quechua:** Aymara-Quechua proportion > 0.70, **Aymara-Quechua-European:** Aymara-Quechua proportion 0.35-0.70, **European:** European proportion > 0.70, **Mapuche-Huilliche:** Mapuche-Huilliche proportion > 0.70, **Mapuche-Huilliche-European:** Mapuche-Huilliche proportion 0.35-0.70, **Other admixture:** Remaining study participants, **Genetic risk of gallstone disease:** Weighted polygenic risk score based on the six risk variants identified for Latin Americans by Joshi et al. and their corresponding summary statistics for Chileans provided by Bustos et al., **FROH:** Sum of runs of homozygosity above 1.5 Mb divided by the total length of the autosomal genome.

**Figure 1** shows the geographical distribution of GBC and GSD odds ratios (ORs, using the Santiago metropolitan region as the reference), BMI and F_ROH_ in the study population. The ratio of GBC and GSD patients was highest in the de los Lagos and de los Ríos regions. Study participants from the de los Ríos region had the highest mean BMI, and F_ROH_ was particularly high in the Araucanía, de los Lagos and de los Ríos regions. Supplementary **Table S1** presents the characteristics of the study participants, who were classified into six categories of genetic ancestry (European: European proportion > 0.70; Aymara-Quechua: Aymara-Quechua proportion > 0.70; Aymara-Quechua-European: Aymara-Quechua proportion 0.35-0.70; Mapuche-Huilliche: Mapuche-Huilliche proportion > 0.70; Mapuche-Huilliche-European: Mapuche-Huilliche proportion 0.35-0.70; Other admixture: Remaining study participants). The Aymara-Quechua group showed the highest median F_ROH_ (0.028, IQR [0.023-0.033]), followed by Mapuche-Huilliche individuals (median F_ROH_ of 0.026, IQR [0.022-0.039]), compared to a median F_ROH_ of 0.007 (IQR [0.005-0.011]) for individuals in the “Other admixture” category.

**Figure 1.**
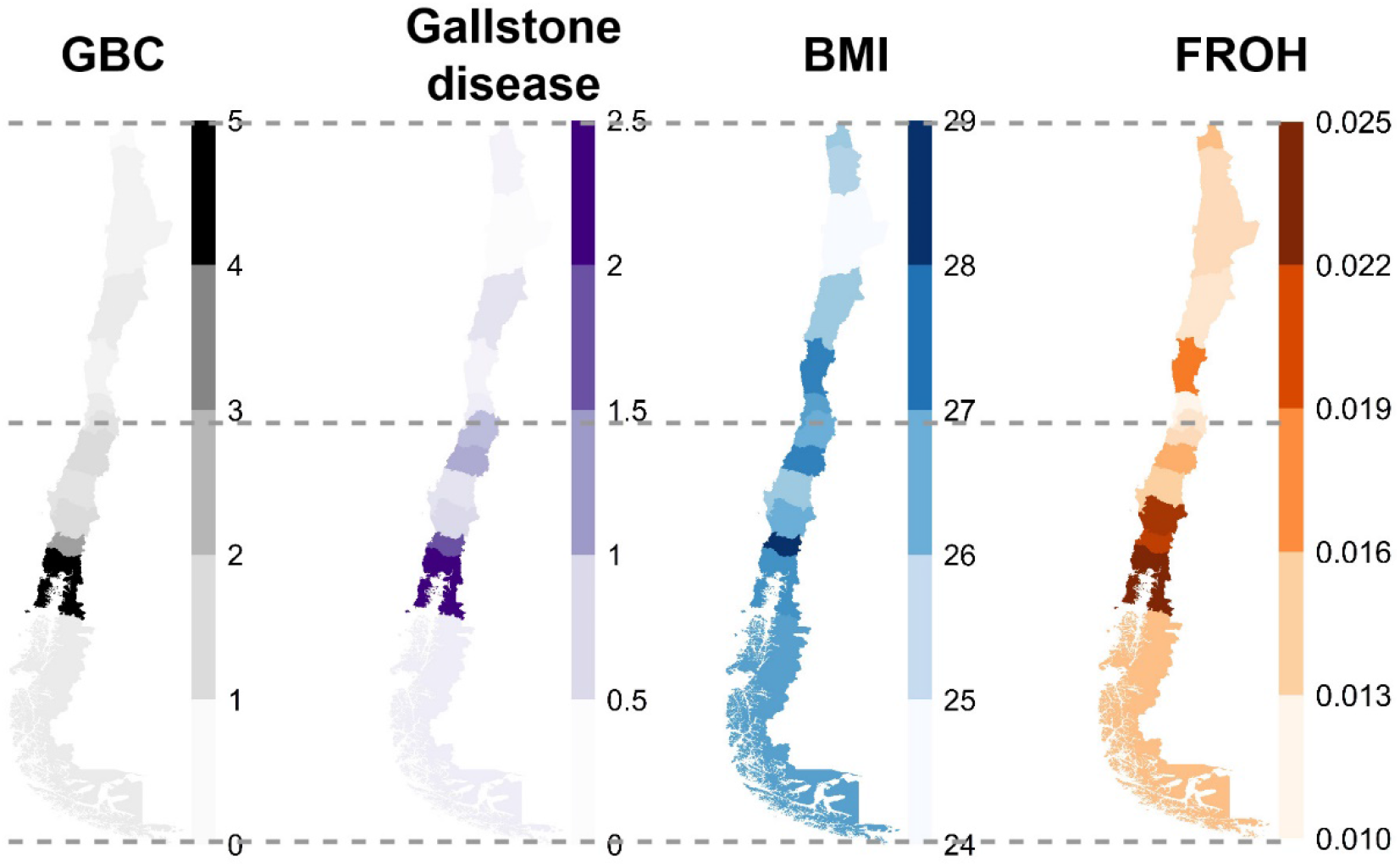
Maps with the distribution of GBC, gallstone disease, BMI and F_ROH_

### Relationship between ROH length and origin, genetic ancestry and GBC risk

ROH size correlates strongly with the time of origin of homozygosity runs. Long ROH indicate a common ancestor a few generations ago, while short ROH point to the shared ancestor being more distant and, consequently, recombination over generations has reduced ROH size. **Figure 2** shows the distribution of ROH size for the five categories of genetic ancestry, and by GBC status. Individuals with a high proportion of indigenous American ancestry exhibited large sums of short ROH (0.3 to 1 Mb) on average, reflecting ancient inbreeding (Aymara-Quechua: 497 Mb ± 52.6, Mapuche-Huilliche: 468 Mb ± 70.1, compared to 230 Mb ± 25.2.x for “Other admixture”; see also Supplementary **Table S1**). Analysis of variance (ANOVA) results confirmed higher total sums of ROH below 1 Mb in both “Aymara-Quechua” and “Mapuche-Huilliche” individuals than in the “Other admixture” category (p-value < 2.6E-16). ROH over 8 Mb represent young autozygous haplotypes that arose less than 5 generations ago and thus reflect cultural practices such as consanguinity, extreme endogamy and/or reproductive isolation. Mapuche-Huilliche individuals had a higher total sum of ROH over 8Mb than the other ancestry categories (ANOVA p-value = 8.2E-13, **Figure 2**). As for the relationship between ROH size and GBC status, neither the differences in the total sum of ROH below 1 Mb, nor the differences in the total sum of ROH above 8 Mb reached the 0.05 statistical significance level.

**Figure 2.**
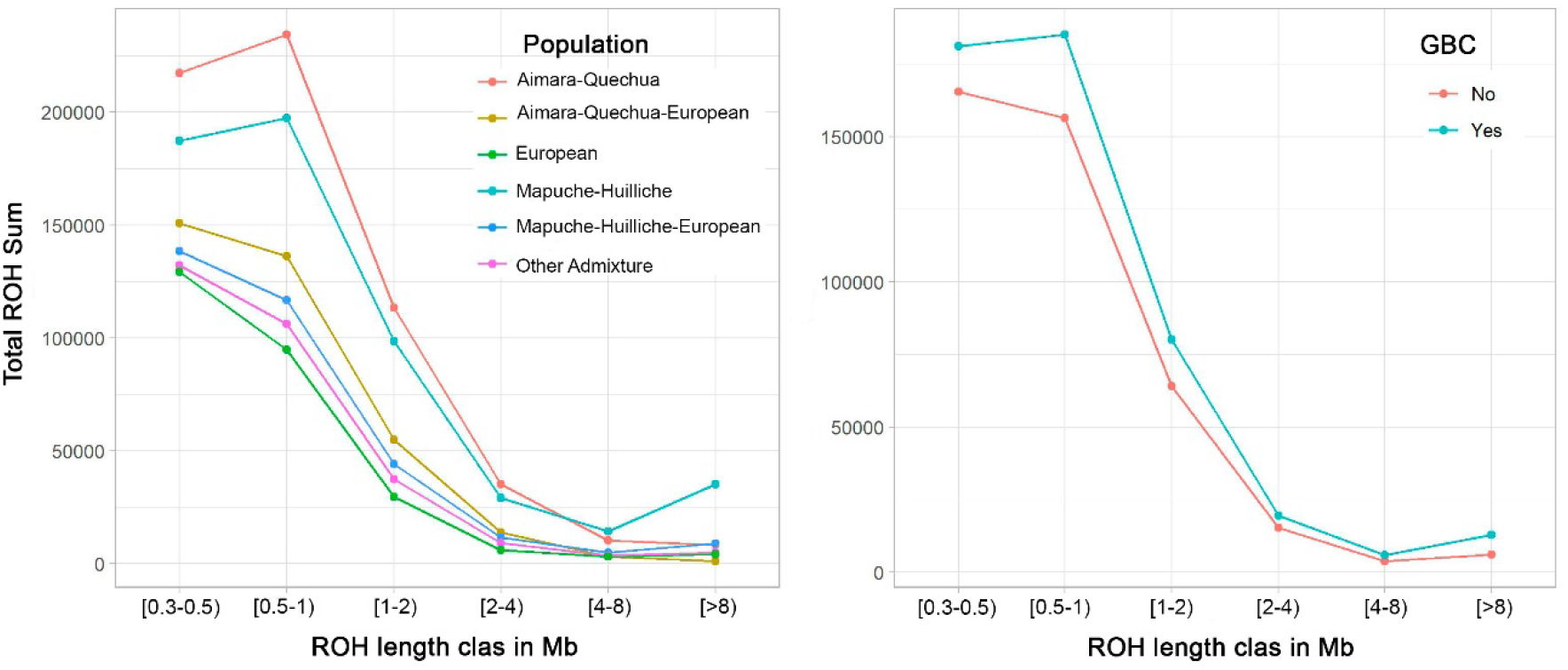
ROH size distribution by population (panel A) and gallbladder cancer (GBC) status (panel B). Represented are ROH total sums over six classes of ROH tract lengths: 0.3≤ROH<0.5 0.5≤ROH<1 Mb, 1≤ROH<2 Mb, 2≤ROH<4 Mb, 4≤ROH<8 Mb and ROH≥8 Mb. Plots are organized by population and presence of GBC. Study individuals were categorized into six groups as follows: **European**: European proportion > 0.70, **Aymara-Quechua**: Aymara-Quechua proportion > 0.70, **Aymara-Quechua-European**: Aymara-Quechua proportion 0.35-0.70, **Mapuche-Huilliche**: Mapuche-Huilliche proportion > 0.70, **Mapuche-Huilliche-European**: Mapuche-Huilliche proportion 0.35-0.70, **Other admixture**: Remaining study participants

We investigated the origin of ROH using two complementary approaches. We examined the relationship between the number and sum of ROH above 1.5Mb, as well as the relationship between F_ROH_ and the systematic inbreeding coefficient (FIS). In the upper panels of **Figure 3**, the relative contributions of genetic drift and consanguinity on homozygosity are examined by comparing the number of ROH (NROH) and the sum of ROH (SROH) per individual genome. When genetic drift is strong, both NROH and SROH are proportionately high. Conversely, consanguinity primarily results in long ROH, leading to a disproportionate increase in SROH compared to NROH. The diagonal lines in the upper panels of **Figure 3** represent the expected relationship between NROH and SROH for an outbred population with no evidence of consanguinity. Individuals with high NROH/SROH values along the diagonal show a high degree of autozygosity caused by genetic drift, while deviations to the right of the diagonal indicate consanguinity. Among the categories of genetic ancestry, especially the “Mapuche-Huilliche”, “Mapuche-Huilliche-European”, and “Other admixed” individuals showed substantial homozygosity attributable to heterogeneous combinations of consanguinity and genetic drift. Comparison with simulated consanguineous mating (**Figure 3**, upper panel left, second cousins in green, first cousins in yellow, avuncular mating (uncle-niece, aunt-nephew, double first cousin) in orange, and incest (brother-sister, parent-offspring) in red) revealed some highly consanguineous individuals in the categories “Mapuche-Huilliche” and “Mapuche-Huilliche-European”. The examination of individuals with and without GBC (**Figure 3**, upper panel right) showed marked heterogeneity within groups, but no notable differences between individuals with/without GBC with regard to their ROH origin.

**Figure 3.**
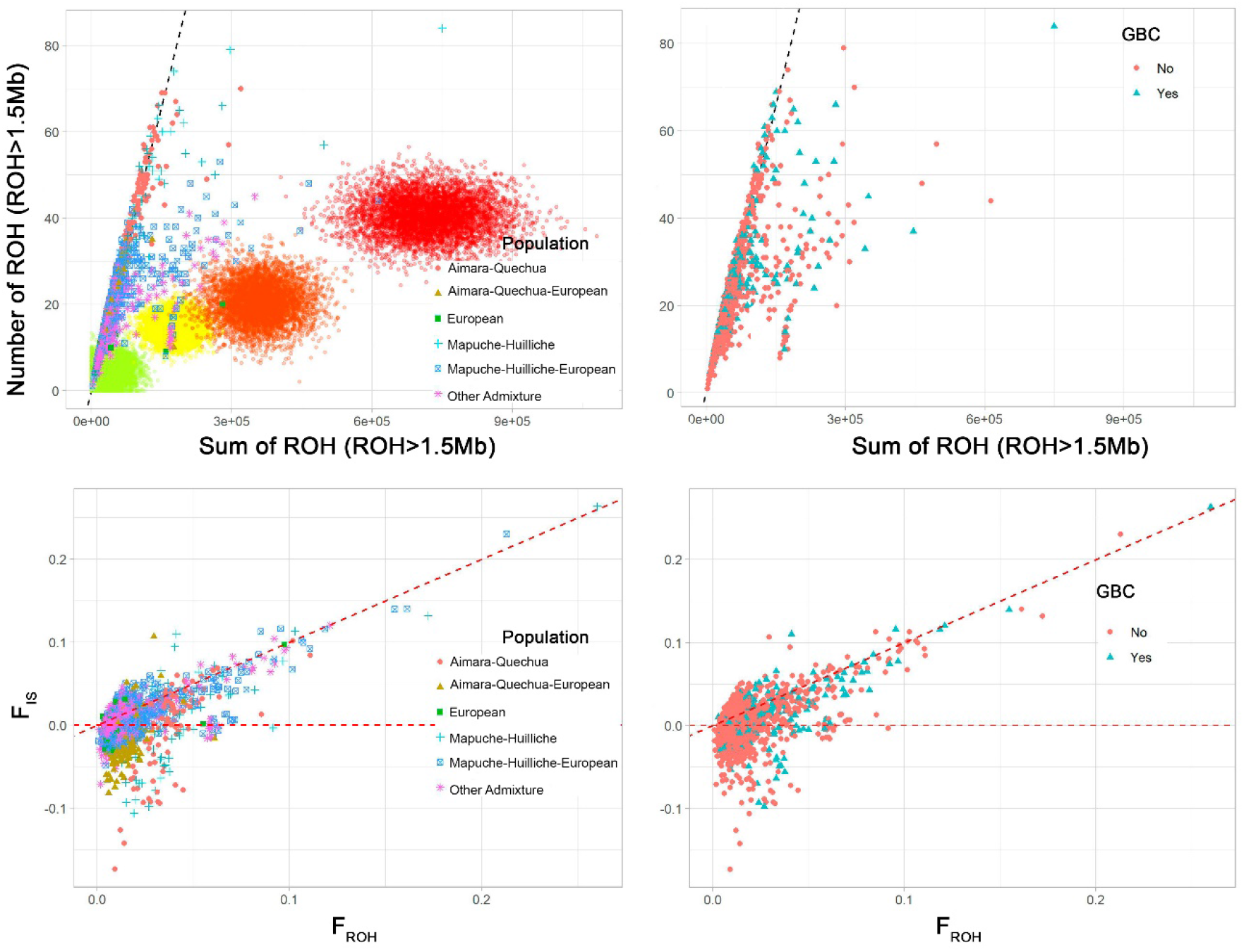
Assessment of ROH origins by population (left panels) and gallbladder cancer (GBC) status (right panels). Study individuals were categorized into six groups as follows: **European**: European proportion > 0.70, **Aymara-Quechua**: Aymara-Quechua proportion > 0.70, **Aymara-Quechua-European**: Aymara-Quechua proportion 0.35-0.70, **Mapuche-Huilliche**: Mapuche-Huilliche proportion > 0.70, **Mapuche-Huilliche-European**: Mapuche-Huilliche proportion 0.35-0.70, **Other admixture**: Remaining study participants. **Upper panels:** Mean number of ROH versus sum of ROH > 1.5Mb for each individual. The dotted straight lines represent the linear regression of the number of ROH on the sum of ROH in individuals of African ancestry in southwestern United States (ASW) and African Caribbean in Barbados (ACB) from the 1000 Genomes Project that represent admixed and thus relatively outbred populations. Simulations of the number and sum of ROH > 1.5Mb for the offspring of different consanguineous mattings are also shown in the left plot. The colour of the dots represents the type of consanguineous mating: second cousin (green), first cousin (yellow), avuncular (uncle-niece, aunt-nephew, double first cousin) (orange), incest (brother-sister, parent-offspring) (red). 5,000 individuals were simulated for each mating type. Note that the simulation did not include drift, but the degree of right shift can be projected to cases where there is a non-zero level of autozygosity due to drift. **Lower panels**: Systematic inbreeding coefficient (F_IS_) versus the FROH-based inbreeding coefficient. F_IS_ represents the average individual single nucleotide polymorphism homozygosity relative to the expected homozygosity of alleles randomly drawn from the population, which was calculated using the -het function in PLINK. The dotted diagonal represents F_IS_ = F_ROH_, and the dotted horizontal line shows F_IS_ = 0.

In the lower panels of **Figure 3**, the mean F_IS_ is plotted against the F_ROH_ for each study participant. The diagonal line (F_IS_ = F_ROH_) and the horizontal line (F_IS_ = 0) delineate three distinct regions. (1) Individuals near the diagonal line have a pronounced component of systematic inbreeding or F_IS_, indicating consanguinity. (2) Individuals near the horizontal line show panmictic inbreeding, caused mainly by genetic drift. (3) Negative F_IS_ values indicate that low effective population size, isolation, and genetic drift play an important role. The lower panels of **Figure 3** show heterogeneity of ROH origin between and within populations, and illustrate that consanguinity plays an important role in the origin of homozygosity in highly inbred individuals. Consistent with the upper panel on the right, differences between individuals with/without GBC in terms of ROH origin are not apparent in the lower right panel.

### Effects of the Homozygosity in the prevalence of GBC

As presented in **Table 2**, statistical analysis confirmed the increased risk of GBC in women, per year (but a decreasing risk per year^2^), in individuals with low levels of education, with increasing proportions of Mapuche-Huilliche ancestry, and with increasing genetic susceptibility for GSD. However, we found no association between F_ROH_ and overall GBC risk. Similarly, no effects of homozygosity on BMI or GSD were observed, as shown in Supplementary **Table S3**. Nevertheless, we identified interaction effects between F_ROH_ and sex, age, and genetic risk of GSD on GBC risk. In light of these intriguing results, we further examined the impact of F_ROH_ after stratifying the complete dataset by sex (Supplementary **Table S4**), age (Supplementary **Table S5**), and genetic risk of GSD (Supplementary **Table S6**).

**Table 2.** Relative risk of gallbladder cancer by potential confounders and FROH.

| Variable | Level | OR | 95% CI | P-value |
| --- | --- | --- | --- | --- |
| Sex | Male | Baseline |  |  |
|  | Female | **3.48** | 2.64 – 4.62 | 2.1E-16 |
| Age | Per year | **1.18** | 1.12 – 1.26 | 1.4E-14 |
| Age<sup>2</sup> | Per year<sup>2</sup> | **0.99** | 0.99 – 1.00 | 2.6E-16 |
| Education | Primary & informal schooling | **2.65** | 1.65 – 4.31 | 1.2E-04 |
|  | Secondary | Baseline |  |  |
|  | Technical | 0.47 | 0.20 – 1.07 |  |
|  | Postgraduate | 0.22 | 0.03 – 1.07 |  |
|  | University | 0.57 | 0.32 – 1.02 |  |
|  | Missing | **0.14** | 0.09 – 0.22 |  |
| BMI | Normal | Baseline |  | 0.35 |
|  | Overweight | 1.34 | 0.89 – 2.04 |  |
|  | Obesity | 1.27 | 0.83 – 1.97 |  |
| Ancestry | Per Aymara-Quechua % | **0.97** | 0.96 – 0.98 | 0.04 |
|  | Per Mapuche-Huilliche % | **1.02** | 1.01 – 1.05 | 1.2E-05 |
| Genetic risk of gallstone disease | Per doubling in disease prevalence | **2.75** | 1.49 – 4.94 | 0.001 |
| FROH | Per 1% | 1.07 | 0.98 – 1.15 | 0.26 |

**OR:** Odds ratio, **CI:** Confidence interval, **P-value:** Probability value, **BMI:** Body mass index, **Normal:** BMI $\leq 25$ kg/m<sup>2</sup>, **Overweight:** BMI 25-29.9 kg/m<sup>2</sup>, **Obesity:** BMI $> 30$ kg/m<sup>2</sup>, **Aymara-Quechua %:** Proportion of northern Chilean Native American ancestry, **Mapuche-Huilliche %:** Proportion of Mapuche-Huilliche ancestry, **Genetic risk of gallstone disease:** Weighted polygenic risk score based on the six risk variants identified for Latin Americans by Joshi et al. and their corresponding summary statistics for Chileans provided by Bustos et al., **FROH:** Sum of runs of homozygosity above 1.5 Mb divided by the total length of the autosomal genome. Bold type indicated that the 95% CI does not include 1.00.

**Figure 4** depicts the ORs from the different analyses conducted. The forest plot illustrates a notable influence of F_ROH_ on GBC risk for specific subsets of the population: males, individuals under 60 years of age (mean age at GBC diagnosis in the study population), and those with a higher than average genetic risk of GSD. Among males, GBC risk increased by 19% for every 1% rise in F_ROH_ (OR = 1.19, 95% CI: 1.01-1.39, p-value = 0.002), but we found no association between F_ROH_ and GBC risk in women. Considering an age cutoff of 60 years (average age of GBC diagnosis), we observed a 30% increase in GBC risk for each 1% increase in F_ROH_ (OR = 1.30, 95% CI: 1.09-1.98), only among individuals younger than 60 years. Stratifying by median genetic risk of GSD, which corresponded to a weighted polygenic risk score of 0.445, individuals with a higher than median genetic susceptibility to GSD showed a 12% increased risk of GBC for every 1% elevation in F_ROH_ (OR = 1.12, 95% CI: 1.03-1.21).

**Figure 4.**
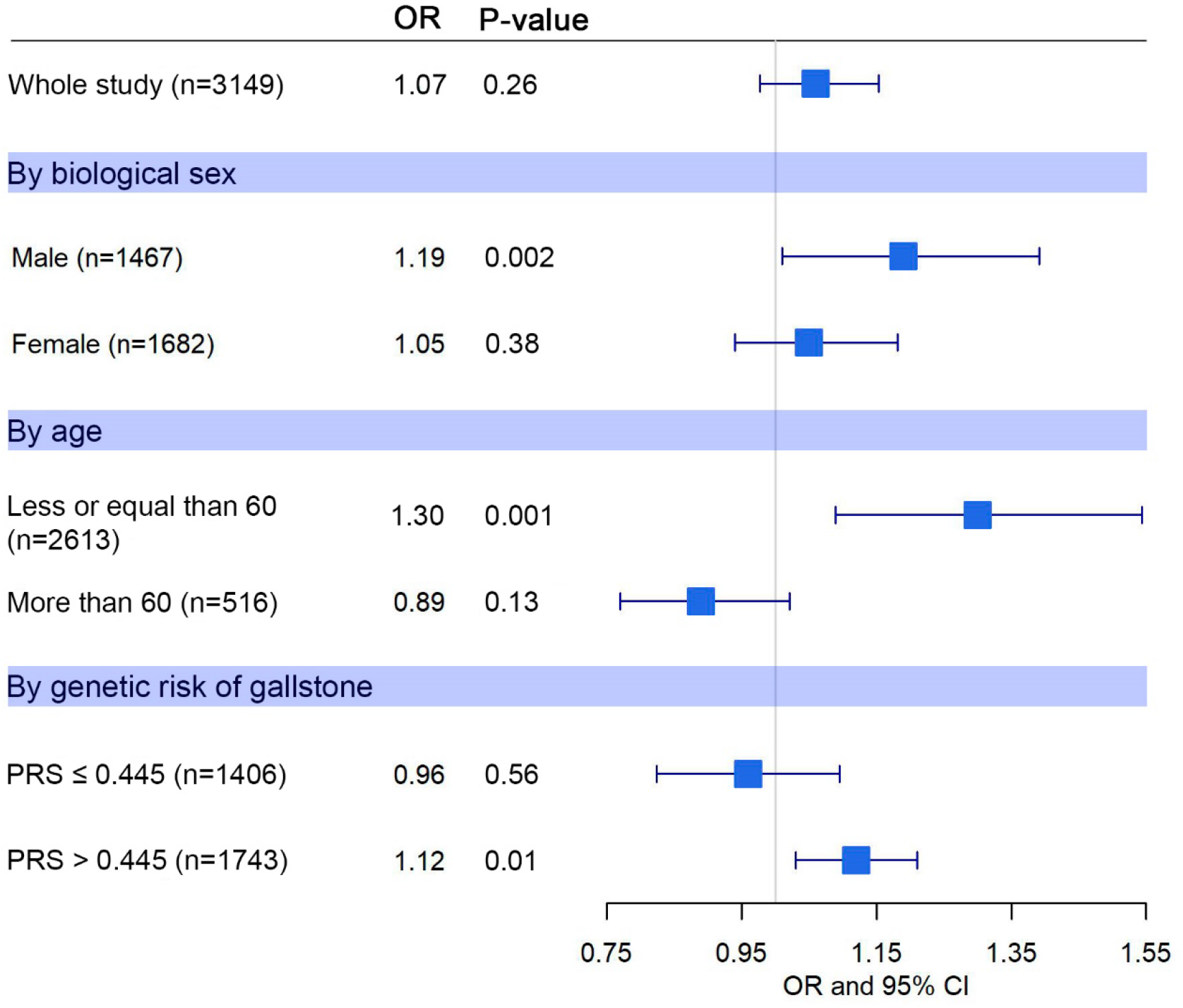
Inbreeding and gallbladder cancer (GBC) risk. Odds ratios (ORs) per 1% FROH with probability values and 95% confidence intervals are shown for the whole study and stratified by biological sex, age (considering a cut-off point of 60 years) and genetic risk of gallstone disease (weighted polygenic risk score (PRS) based on the six risk variants identified for Latin Americans by Joshi et al. and their corresponding summary statistics for Chileans provided by Bustos et al., considering the median score of 0.445 as cut-off point).

## Discussion

GBC continues to pose a significant challenge to the healthcare system in high incidence areas due to very limited treatment options for advance disease, and the absence of early detection markers (18). It has been postulated that GBC takes 10-20 years to develop, typically following the sequence of gallstones and inflammation, gallbladder dysplasia and GBC, and that surgical removal of the gallbladder (cholecystectomy) is an effective option for prevention before the onset of symptoms, emphasizing the urgent need to identify and exploit risk and early diagnosis factors associated with this malignancy. The highly variable prevalence of GBC in different subpopulations and geographic regions, as well as the familial aggregation of GBC (19)suggest a genetic component to GBC risk. Among the large differences in prevalence, GBC is the third leading cause of death in Japanese living in the United States and the third leading malignancy in the Native American population, according to the New Mexico Tumor Registry (https://hsc.unm.edu/new-mexico-tumor-registry/ last checked 26 February 2024). Conversely, GBC appears to be rare in people of African descent. Importantly for this study, clear associations have been reported between Asian and indigenous American ancestries, and increased susceptibility to GBC. However, even within these broad ethnic groups, the distribution of GBC is very heterogeneous.

Inbreeding has been associated with GBC risk in the past. For example, the Abiquiu community in the Chama Valley has both a high prevalence of GBC and endogamous mating practices that have led to high levels of inbreeding, suggesting a potential link between homozygosity and GBC susceptibility (4). In Chile, one of the countries with the highest GBC incidence in the world, the individual proportion of overall indigenous American ancestry does not correlate with GBC mortality, but the specific indigenous Mapuche subcomponent (the Mapuche are the largest indigenous people living mainly in central and southern Chile) is strongly associated with GBC incidence and mortality. Considering this scenario, we investigated the genetic contribution to GBC risk from a new perspective —assessing the potential influence of ancient and recent inbreeding quantified by the genomic distribution of ROH. Our study is the first attempt to examine the relationship between GBC, homozygosity (quantified as the fraction of the genome in ROH over 1.5 Mb), and the proportion of indigenous American ancestry present in Chile. Of note, homozygosity exhibited a considerable degree of variability across the six categories of genetic ancestry defined in the present study, which is consistent with previous large-scale investigations.

Our study provides novel insights into the interplay of genetic ancestry, homozygosity and GBC development. The particular genetic tapestry of Chile, woven through a complex history of admixture and migration, provides an optimal framework for such studies. The six defined ancestry categories exhibited different characteristics in terms of ROH, mirroring their unique genetic history. This variability translates into improved statistical power, which distinguishes our study from analyses based on European cohorts. Remarkably, the groups with indigenous American ancestry, in particular Aymara-Quechua individuals, displayed larger average ROH sizes, which can be attributed to ancient inbreeding. In contrast, the presence of longer ROH in the Mapuche-Huilliche category points to consanguinity, shedding light on the diverse origins of homozygosity in these populations.

The crux of our study was to investigate the impact of genomic homozygosity, quantified through F_ROH_, on GBC risk. We simultaneously considered F_ROH_, the proportion of Aymara–Quechua and Mapuche–Huilliche ancestry, as well as BMI, genetic risk of GSD, and education level using logistic regression to assess the effect of homozygosity on GBC risk while accounting for potential confounders. The relevance of considering potential cultural and social confounding, as we did in our study by accounting for educational attainment and individual ancestry proportions, was well illustrated in a comprehensive meta-analysis that scrutinized full-sibling data. Remarkably, F_ROH_ differences between siblings were solely due to Mendelian segregation and remained unaffected by cultural and socioeconomic influences. On average, F_ROH_ effect estimates derived from sibling relationships were 22% lower than their population-based counterparts for all traits analyses, possibly reflecting the contribution of non-genetic confounders.

In contrast to comparisons between separate ethnic groups (e.g., individuals of European versus Mapuche ancestry), our study relied on data from genetically admixed Chileans with continuous gradients of homozygosity and ancestry, which lent robustness to our findings by attenuating the influence of sociocultural confounders. Although no overarching association emerged across the entire dataset, we were able to unveil strong interaction effects between F_ROH_ and sex, age, and genetic risk of GSD. Intriguingly, the results suggested a notable influence of F_ROH_ on the development of GBC in certain population group, particularly men, individuals under 60 years of age (men and women), and those with genetic predisposition to gallstones. Notably, the absence of a F_ROH_ effect in women points to intricate gender differences in GBC development. We found no interaction between the Mapuche-Huilliche subcomponent of indigenous American ancestry and F_ROH_, suggesting that inbreeding affects GBC risk independent of genetic ancestry.

In conclusion, the present study indicates a complex interplay between F_ROH_ and GBC risk, pointing to stronger inbreeding effects in men, individuals younger than 60 years, and persons with an increased genetic risk of GSD. Replication of these results in an independent cohort, ideally with a larger study population and including additional sociocultural covariates, would undoubtedly underpin the robustness of our findings. The results indicate that Mapuche-Huilliche ancestry and inbreeding act as independent determinants of genetic susceptibility to GBC, which is important from both a scientific and a preventive perspective. Our study contributes to a deeper understanding of the multifaceted factors underlying the development of GBC, and sets the stage for further investigation of the complex interplay between homozygosity, genetic ancestry, and disease susceptibility.

## Materials and Methods

### Study population and ethics approvals

The phenotype and genotype data analysed in this study has been used previously to investigate the relationship between indigenous American ancestry, GBC, GSD and BMI (12). The present study included 202 additional GBC and 582 additional GSD patients recruited according to a study protocol that complied with the ethical guidelines of the 1975 Declaration of Helsinki, and was approved by the ethics committees of Servicio de Salud Metropolitano Oriente, Santiago de Chile (#06.10.2015, #08.03.2016 and #12.11.2019), Servicio de Salud Metropolitano Sur Oriente, Santiago de Chile (#15.10.2015 and #05.04.2018), Servicio de Salud Metropolitano Central, Santiago de Chile (#1188-2015), Servicio de Salud Coquimbo, Coquimbo, Chile (#01.04.2016), Servicio de Salud Maule, Talca, Chile (#05.11.2015), Universidad Católica del Maule, Talca, Chile (#102-2020), Servicio de Salud Concepción, Concepción, Chile (ID: 16-11-97 and ID:19-12-111), Servicio de Salud Araucanía Sur, Temuco, Chile (#10.02.2020), Servicio de Salud Valdivia, Valdivia, Chile (ID:438), Centro de Bioética, Universidad del Desarrollo, Clínica Alemana de Santiago, Santiago de Chile (#2018-97, ID 678) and Unidad de Investigación Hospital San Juan de Dios, Santiago de Chile (#6182), the Medical Faculties of Universidad de Chile (approval #123-2012 and #11.10.2012) and Pontificia Universidad Católica de Chile (#11-159). In 77% of GBC patients, the diagnosis was made after surgical removal of the gallbladder (cholecystectomy), and gallstones were found in around 86% of the GBC patients investigated. GSD patients were patients who underwent cholecystectomy for symptomatic gallstones. The remaining study participants belonged to population-based studies with a BMI distribution that was representative of the general Chilean population (12).

All participants provided written informed consent prior to enrolment in in the study, using a consent form reviewed by a representative of the [Chilean Foundation of Gastrointestinal Cancer Patients](http://www.gist.cl/). This representative is also a permanent member of the External Advisory Board of the European-Latin American Consortium towards Eradication of Preventable Gallbladder Cancer – [EULAT Eradicate GBC](http://www.saludvesiculabiliar.org/), which meets annually to discuss the project objectives, progress and relevance of the project results to patients. The EULAT Eradicate GBC dissemination videos are available in Aymara, Quechua and Mapudungun, the language of the Mapuche people. To improve the communication of study results related to ancestry, we have organized a <u>Symposium at the joint meeting of the Chilean Genetics Society and the Chilean Society of Evolution</u>, and recently held a [Summer School on ancestry and molecular health](https://www.klinikum.uni-heidelberg.de/medizinische-biometrie/forschung/arbeitsgruppen/statistical-genetics/research-training).

### ROH calling

ROH longer than 300 Kb were called using PLINK v1.9 software (20) and the following parameters: -- *homozyg-snp 30* (minimum number of single nucleotide polymorphisms (SNPs) a ROH must have), -- *homozyg-kb 300* (length of sliding window in Kb), --*homozyg-density 30* (minimum density required to consider a ROH, 1 SNP in 30 Kb), --*homozyg-window-snp 30* (number of SNPs the sliding window must have), --*homozyg-gap 1000* (length in Kb between two SNPs to be considered in two different segments), --*homozyg-window-het 1* (number of heterozygous SNPs allowed in a window), --*homozyg-window-missing 5* (number of missing calls allowed in a window), --*homozyg-window-threshold 0*.*05* (proportion of the overlapping window that must be called homozygous to define a given SNP as “in a homozygous segment”). No linkage disequilibrium pruning was performed. We filtered out SNPs with minor allele frequencies <0.01 and those deviating from Hardy-Weinberg (H-W) proportions with a p-value <0.001. These parameters have already been used and validated in large-scale, published studies, and they have been shown to call ROH corresponding to autozygous segments in which all SNPs (including those not present on the genotyping array) are homozygous-by-descent (13, 15).

### Estimating inbreeding and its origin

Inbreeding can arise from departure from panmixia, which involves systematic inbreeding, also known as consanguinity (F_IS_), or from genetic isolation and a small effective population size, genetic drift (F_ST_), which leads to panmictic inbreeding (21, 22). Systematic inbreeding directly affects the H-W equilibrium of a population, but its effects can be reversed within a single generation of panmictic breeding. In contrast, panmictic inbreeding does not affect H-W proportions, but leads to a reduction in genetic variability within the population though allele loss (23). The total inbreeding coefficient F_IT_ is defined as the probability that an individual receives two alleles identical-by-descent: (1-F_IT_) = (1-F_IS_)(1-F_ST_) (24, 25). Traditionally, F_IT_ has been measured using deep genealogies. Here we considered F_ROH_, or the genomic inbreeding coefficient, as a proxy for F_IT_, and estimated F_IS_ using SNP data.

F_IS_ is the average SNP homozygosity within an individual relative to the expected homozygosity of alleles randomly drawn from the population. PLINK estimates F_IS_ using the following expression:

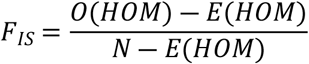

Where *Observed Hom* is the observed number of homozygous SNPs, *Expected Hom* is the expected number of homozygous SNPs considering H–W proportions, and *N* is the total number of non-missing genotyped SNPs. F_IS_ thus measures inbreeding in the current generation, with F_IS_ = 0 indicating random mating, F_IS_ > 0 indicating consanguinity and F_IS_ < 0 indicating inbreeding avoidance.

F_ROH_ quantifies the actual proportion of the autosomal genome that is autozygous over and above a specific minimum length ROH threshold. When analysing ROH>1.5Mb, F_ROH_ correlates strongly (r=0.86) with inbreeding coefficients obtained from six-generation pedigrees (26).

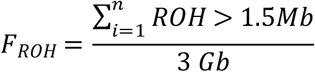

### Testing inbreeding depression

Traditionally, inbreeding depression refers to the decline in the evolutionary fitness of an individual or population due to an increase in homozygosity as a result of inbreeding. This concept has now been extended to any complex trait, describing the change in average phenotypic value within a population due to inbreeding. When considering the combined influence of all loci affecting a specific trait, in terms of the additive combination of genotypic values, the average trait value within a population with an inbreeding coefficient (F) is given by (27):

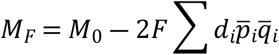

Here, M_0_ stands for the average population value prior to inbreeding, d is the genotypic value of heterozygotes, and p and q denote the allele frequencies.

This equation illustrates that inbreeding leads to a change in the average trait value within a population when the cumulative genotypic value of heterozygotes (d) is not zero, indicating that the trait must exhibit some form of directional dominance or overdominance in its genetic architecture. Furthermore, for additive locus combinations, the change in the mean due to inbreeding is directly proportional to the inbreeding coefficient (28). This knowledge enables to identify instances of inbreeding depression in complex traits showing directional dominance through regression analysis, provided that the population under study has a certain degree of inbreeding. It is important to note that the underlying genetic architecture of a trait, including the effects of inbreeding depression, may be different in different populations. The severity of inbreeding depression and the genetic basis of a trait depend on factors such as selection pressure, environmental influences, and population structure, which lead to variations in genetic frequencies between populations.

In this study, we assessed the relationship between F_ROH_ and GBC risk using multiple logistic regression. GBC status was regressed against F_ROH_ as an independent variable, along with age, age^2^, biological sex, education, proportions of Aymara-Quechua and Mapuche-Huilliche ancestry, BMI, and genetic risk of GSD disease, characterised by a weighted polygenic risk score based on six GSD-associated variants previously proposed for the Chilean population (29). The interactions of F_ROH_ and sex, age, genetic risk of GSD and ancestry proportions were also tested.

## Data Availability Statement

Files with the called ROHs cannot be made publicly available due to privacy and ethical restrictions (potential de-identification of study participants), but they are available on request from the corresponding author. The source code to reproduce all the results described is provided as supplementary material and available at www.biometrie.uni-heidelberg.de/StatisticalGenetics/Software_and_Data.

## Acknowledgments

The authors gratefully acknowledge the data storage service SDS@hd supported by the Ministry of Science, Research, and the Arts Baden-Württemberg (MWK) and the German Research Foundation (DFG) through grants INST 35/1314-1 FUGG and INST 35/1503-1 FUGG.

## References

1. Eslick GD. Epidemiology of gallbladder cancer. Gastroenterol Clin North Am. 2010;39(2):307–30, ix.

2. Srivastava K, Srivastava A, Kumar A, Mittal B. Gallbladder cancer predisposition: a multigenic approach to DNA-repair, apoptotic and inflammatory pathway genes. PLoS One. 2011;6(1):e16449.

3. Salazar M, Ituarte C, Abriata MG, Santoro F, Arroyo G. Gallbladder cancer in South America: epidemiology and prevention. Chin Clin Oncol. 2019;8(4):32.

4. Devor EJ, Buechley RW. Gallbladder cancer in Hispanic New Mexicans: I. General population, 1957-1977. Cancer. 1980;45(7):1705–12.

5. Lorenzo Bermejo J, Boekstegers F, Gonzalez Silos R, Marcelain K, Baez Benavides P, Barahona Ponce C, et al. Subtypes of Native American ancestry and leading causes of death: Mapuche ancestry-specific associations with gallbladder cancer risk in Chile. PLoS Genet. 2017;13(5):e1006756.

6. Andia ME, Hsing AW, Andreotti G, Ferreccio C. Geographic variation of gallbladder cancer mortality and risk factors in Chile: a population-based ecologic study. Int J Cancer. 2008;123(6):1411–6.

7. Bertran E, Heise K, Andia ME, Ferreccio C. Gallbladder cancer: incidence and survival in a high-risk area of Chile. Int J Cancer. 2010;127(10):2446–54.

8. Nogueira L, Foerster C, Groopman J, Egner P, Koshiol J, Ferreccio C, et al. Association of aflatoxin with gallbladder cancer in Chile. JAMA. 2015;313(20):2075–7.

9. Reich D, Patterson N, Campbell D, Tandon A, Mazieres S, Ray N, et al. Reconstructing Native American population history. Nature. 2012;488(7411):370–4.

10. Ruiz-Linares A, Adhikari K, Acuna-Alonzo V, Quinto-Sanchez M, Jaramillo C, Arias W, et al. Admixture in Latin America: geographic structure, phenotypic diversity and self-perception of ancestry based on 7,342 individuals. PLoS Genet. 2014;10(9):e1004572.

11. Eyheramendy S, Martinez FI, Manevy F, Vial C, Repetto GM. Genetic structure characterization of Chileans reflects historical immigration patterns. Nat Commun. 2015;6:6472.

12. Zollner L, Boekstegers F, Barahona Ponce C, Scherer D, Marcelain K, Garate-Calderon V, et al. Gallbladder Cancer Risk and Indigenous South American Mapuche Ancestry: Instrumental Variable Analysis Using Ancestry-Informative Markers. Cancers (Basel). 2023;15(16).

13. Ceballos FC, Joshi PK, Clark DW, Ramsay M, Wilson JF. Runs of homozygosity: windows into population history and trait architecture. Nature Reviews Genetics. 2018;19(4):220–34.

14. Ceballos FC, Hazelhurst S, Clark DW, Agongo G, Asiki G, Boua PR, et al. Autozygosity influences cardiometabolic disease-associated traits in the AWI-Gen sub-Saharan African study. Nat Commun. 2020;11(1):5754.

15. Clark DW, Okada Y, Moore KHS, Mason D, Pirastu N, Gandin I, et al. Associations of autozygosity with a broad range of human phenotypes. Nature Communications. 2019;10(1):4957.

16. Mooney JA, Huber CD, Service S, Sul JH, Marsden CD, Zhang Z, et al. Understanding the Hidden Complexity of Latin American Population Isolates. Am J Hum Genet. 2018;103(5):707–26.

17. Koenigstein F, Boekstegers F, Wilson JF, Fuentes-Guajardo M, Gonzalez-Jose R, Bedoya G, et al. Inbreeding, Native American ancestry and child mortality: linking human selection and paediatric medicine. Hum Mol Genet. 2022;31(6):975–84.

18. Henley SJ, Weir HK, Jim MA, Watson M, Richardson LC. Gallbladder Cancer Incidence and Mortality, United States 1999-2011. Cancer Epidemiol Biomarkers Prev. 2015;24(9):1319–26.

19. Hemminki K, Hemminki A, Forsti A, Sundquist K, Li X. Genetics of gallbladder cancer. Lancet Oncol. 2017;18(6):e296.

20. Purcell S, Neale B, Todd-Brown K, Thomas L, Ferreira MAR, Bender D, et al. PLINK: A tool set for whole-genome association and population-based linkage analyses. American Journal of Human Genetics. 2007;81(3):559–75.

21. Jacquard A. Inbreeding - One word, several meanings. Theoretical Population Biology. 1975;7(3):338–63.

22. Templeton A R, Read B. Inbreeding, One Word, Several Meanings, Much Confusion. Biological Conservation. 1996;75(311).

23. Hedrick PW. Genetics of populations. sudbury: Jones and Bartlett Publishers; 2005.

24. Hartl DL, Clark AG. Principles of population Genetics. Sunderland: Sinauer Associates; 2007.

25. Weir BS. Estimating F-statistics: A historical view. The British Journal for the Philosophy of Science. 2012;79(5):637–43.

26. McQuillan R, Leutenegger AL, Abdel-Rahman R, Franklin CS, Pericic M, Barac-Lauc L, et al. Runs of homozygosity in European populations. American Journal of Human Genetics. 2008;83(3):359–72.

27. Falconer DS, Mackay TFC. Quantitative genetics 4 edition ed: Pearson; 1996.

28. Charlesworth D, Willis JH. The genetics of inbreeding depression. Nat Rev Genet. 2009;10(11):783–96.

29. Barahona Ponce C, Scherer D, Brinster R, Boekstegers F, Marcelain K, Garate-Calderon V, et al. Gallstones, Body Mass Index, C-Reactive Protein, and Gallbladder Cancer: Mendelian Randomization Analysis of Chilean and European Genotype Data. Hepatology. 2021;73(5):1783–96.

